# Orchid mycorrhizal communities associated with *Orchis italica* are shaped by ecological factors and geographical gradients

**DOI:** 10.1101/2024.07.06.601878

**Authors:** Marco G. Balducci, Jacopo Calevo, Karl J. Duffy

## Abstract

**Aim:** The influence of mutualists on plant distributions is only beginning to be understood. Orchids depend on orchid mycorrhizal (OrM) fungi to germinate, yet the distribution of OrM and how they vary according to both abiotic and biotic factors is unclear. We investigated the abundance and diversity of OrM communities associated with the Mediterranean orchid *Orchis italica* and quantified how they vary according to both geographical and ecological factors.

**Location:** Mediterranean Basin.

**Taxon:** *Orchis italica* Poir. (Orchidaceae)

**Methods:** We used metabarcoding of the ITS2 region to identify OrM fungi associated with adult individuals in 23 populations of *O. italica* across latitudinal and longitudinal gradients in the Mediterranean region. We used both multivariate analyses and Joint Species Distribution Models (JSDMs) based on geographical, climate, and soil variables to test how both common OrM fungi and their communities vary according to geographical and ecological factors.

**Results:** Eighty OrM OTUs were found associating with *O. italica*. However, five Tulasnellaceae OTUs and one Ceratobasidiaceae OTU were found in every population. Abundance of these taxa, as measured by number of reads, increased from west to east and decreased from south to north, indicating OrM abundance may be determined by geographical gradients. OrM community composition varied according to precipitation, annual mean temperature, and soil phosphorous content. JSDMs revealed there were both positive and negative co-occurrences among these ubiquitous OrM.

**Main Conclusions:** Despite associating with many OrM across its range, only six OrM were widespread, indicating that *O. italica* may be an apparent generalist in its association with OrM. Abundance of these OrM is determined by geographical gradients and ecological factors. This highlights the importance of quantifying the distribution of belowground mutualists in understanding the limits to plant distributions.

## Introduction

Plants often depend on mutualists to complete their life cycle, which can influence both their demography and range limits (Lau et al., 2008; Sexton et al., 2009). Mutualisms contribute to ecosystem functioning and have shaped much of global biodiversity (Chomicki et al., 2019; Schemske et al., 2009; Tedersoo et al., 2020). One of the most common and widespread interactions is the mutualistic relationship between plants and mycorrhizal fungi (Brundrett, 2009; Brundrett & Tedersoo, 2018). Approximately 90% of vascular plants form associations with mycorrhizal fungi (van Der Heijden et al., 2015; Smith & Read, 2010). Consequently, mycorrhizal symbioses play important roles in determining individual plant fitness and plant community composition (van Der Heijden et al., 2015). Understanding large-scale patterns of mycorrhizal diversity can help explain global patterns in plant diversity and can inform predictions and mitigate ecosystem responses to ongoing global changes (Tedersoo et al., 2020). Yet, despite their importance, the geographical distributions and ecology of mycorrhizas, are poorly understood (Kõljalg et al., 2013; Tedersoo et al., 2014; Van Der Putten, 2017).

Members of the Orchidaceae obligately depend on mycorrhizal fungi, the orchid mycorrhizal fungi (OrM), for seed germination and subsequent establishment (Dearnaley et al., 2016). Hence, the availability and abundance of fungal symbionts may restrict their distributions (McCormick et al., 2018). Most photosynthetic orchids form mycorrhizas with fungi from one or more of the fungal families Ceratobasidiaceae, Sebacinaceae, Serendipitaceae, and Tulasnellaceae (Dearnaley et al., 2012; Weiß et al., 2016). Several studies have shown that orchids generally do not associate with one single fungal taxon, but with multiple OrM simultaneously, often from different families (Jacquemyn et al., 2015; Waterman et al., 2011; Waud et al., 2016). The number and composition of fungal taxa associating with a single orchid species is variable, and detection of OrM may depend on orchid developmental stage (Bidartondo & Read, 2008; Waud et al., 2017) and ecological conditions in which the orchid occurs (Duffy et al., 2019; Jacquemyn et al., 2016). Although mycorrhizal dependency is known to be an important factor influencing both the distribution and abundance of orchid populations (McCormick et al., 2018), few studies have quantified the relative importance of environmental factors in driving OrM fungi at large spatial scales.

Over large spatial scales, variation in factors such as climate, latitude, and edaphic conditions may explain soil fungal diversity (Tedersoo et al., 2014; Treseder et al., 2014). However, abiotic variables that can explain the distribution of orchid individuals and populations is generally not understood (Jacquemyn et al., 2017). Since OrM fungi are endophytes and often difficult to detect outside of host roots, we often rely on molecular methods, such as metabarcoding of orchid roots or direct isolation of fungi from roots, to determine which OrM fungi are present on an individual plant. The few available studies that have examined the influence of abiotic conditions as potential drivers of OrM fungal distribution patterns showed that soil moisture content, pH and organic phosphorous were the most important factors determining their occurrence (McCormick & Jacquemyn, 2014; Vogt-Schilb et al., 2020; Waud et al., 2017). A recent broad-scale survey across multiple populations of *Spiranthes spiralis* showed that the diversity of OrM fungal communities is highly variable between habitats and can depend on both spatial and environmental conditions (Duffy et al., 2019). Overall, these studies show that differences in habitat conditions can affect the occurrence of particular OrM fungi and therefore have an impact on plant-fungus interactions in orchids. However, since orchid species are often associated with a particular subset of OrM fungi, it can be difficult to understand the biogeographic patterns of these OrM taxa.

The genus *Orchis* L., contains 21 species that mainly occur in Europe, as well the Middle East and North Africa (Kretzschmar et al., 2007). Previous molecular studies describing the OrM fungal communities associated with the genus revealed a potential broad specificity, with species primarily associating with several fungal Operational Taxonomic Units (OTUs) of the Tulasnellaceae (Girlanda et al., 2011; Jacquemyn et al., 2010, 2011; Mennicken et al., 2023; Oja et al., 2017; Shefferson et al., 2008). Isolation from adult roots of *O*. *italica*, a widespread Mediterranean species, revealed a relatively narrow specialization on members of *Tulasnella* for germination and early development (Balducci et al., 2024). However, our understanding of the whole fungal community in roots over large spatial scales and how they might influence the distribution of *O*. *italica* remains largely unknown. In this study, we tested whether, (1) OrM fungal community composition and abundance of common OrM fungi varies according to latitude and longitude, (2) abiotic factors influence OrM fungal community composition, and (3) whether taxa respond to key environmental variables similarly and whether there are positive and negative residual co-occurrences beyond variation explained by environmental variables.

## Materials and methods

### Study species

*Orchis italica* Poir. is a terrestrial orchid confined in the Mediterranean region. Plants typically grow in open grassland habitats on calcareous soils (Kretzschmar *et al*., 2007). It is a nonrewarding species that is pollinated by a wide range of generalist pollinators that are attracted to the nectarless but brightly coloured flowers. Flowering occurs from April until the beginning of June.

### Sampling

Individual *O. italica* plants from 23 populations were sampled across a 550 km latitudinal range, and a 950 km longitudinal range in 2021 and 2022 (Fig. 1). To identify the OrM fungi, the roots of 10 haphazardly selected individuals were collected from each population, ensuring that minimal damage was caused. After the roots had been excavated from the soil, they were stored in ziplock plastic bags at 4 °C until further processing. Roots were surface sterilized for 3 mins in 1% sodium hypochlorite, washed in sterile distilled water and microscopically checked for fungal colonization. Up to five roots per population were either processed immediately or stored at - 20°C prior to molecular analyses.

**Fig. 1.**
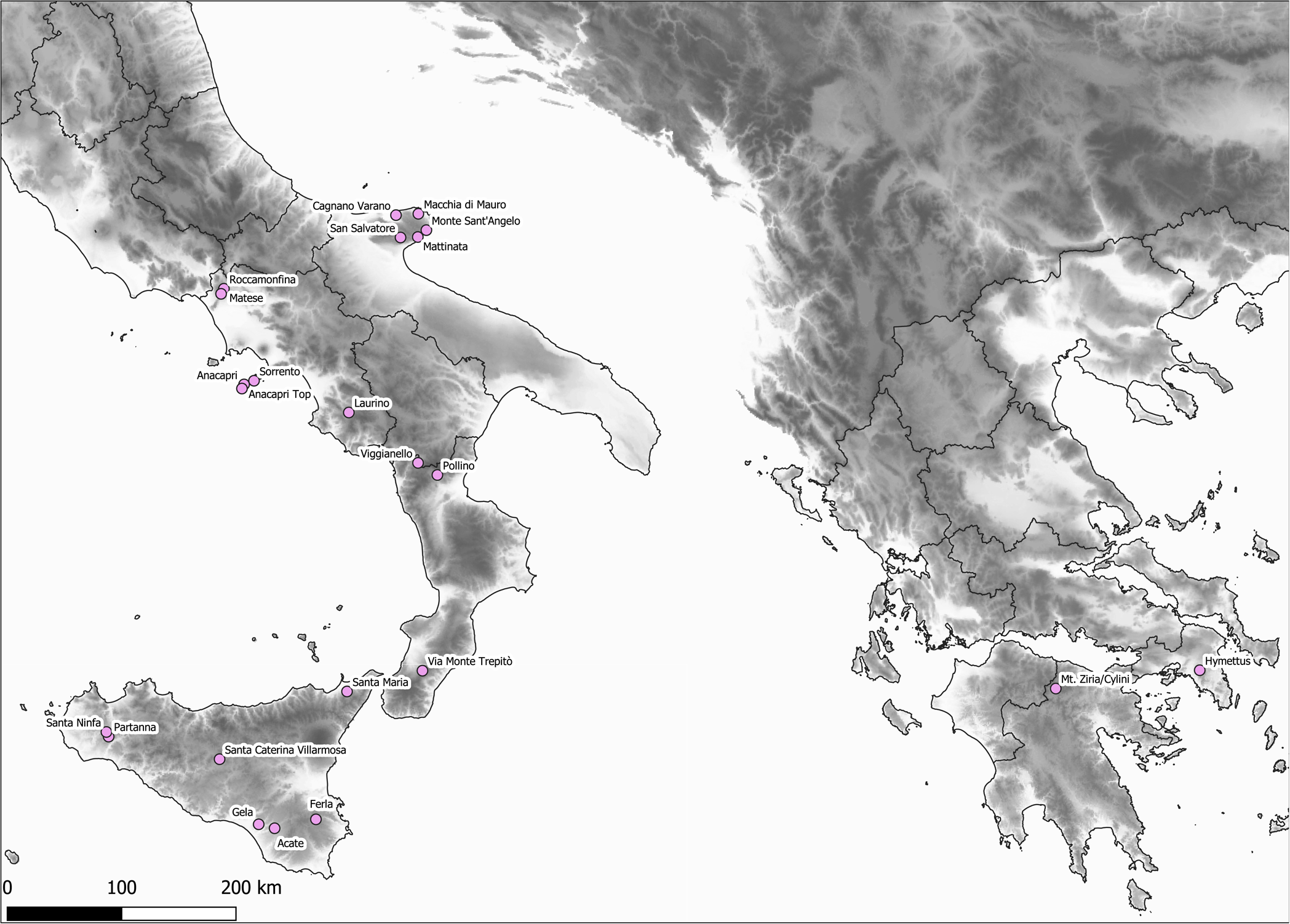
Distribution of the 23 sampled populations of *Orchis italica*.

### Molecular analyses

Genomic DNA was extracted from 0.25 g of highly colonised root fragments using the DNeasy Plant Mini Kit (Qiagen, Hilden, Germany) following the manufacturer’s protocol. The quantity and purity of DNA extractions were checked using a NanoDrop 2000 Spectrophotometer (Thermo Fisher Scientific, Wilmington, DE, USA). Amplifications were performed by means of a semi-nested approach. In the first PCR step, the internal transcribed region (ITS) was amplified using three sets of primers; (i) ITS1-OFa/ITS-OFb/ITS4-OF (Taylor & McCormick, 2008), (ii) ITS1/ITS4tul (Taylor & McCormick, 2008), and (iii) a pair specifically designed for Tulasnellaceae OrM fungi, 5.8S-OF/ITS4Tul (Vogt-Schilb et al., 2020). The PCR amplification was performed for these primer combinations in 25 μl reaction volumes containing 12.5 μl Taq Polymerase (DreamTaq Green, Thermo Scientific, Waltham, USA), 0.5 μl of bovine serum albumin 0.4% (w/v), 1 μl of 0.5μM forward primer, 1 μl 0.5μM reverse primer, 1 μL of genomic DNA (20 ng/μL) and 9 μl nuclease free water. PCR conditions for each primer combination were; (1) initial denaturation of 2 min at 96 °C followed by 35 cycles of 30 s at 94 °C, 40 s at 58 °C and 45 s at 72 °C; (2) initial denaturation of 5 min at 95 °C, followed by 35 cycles of 30 s at 94 °C, 45 s at 54 °C and 1 min at 72 °C and (3) initial denaturation of 30 s at 98 °C, followed by 30 cycles of 10 s at 98 °C, 30 s at 55 °C and 15 s at 72 °C. A further PCR step was then performed on each PCR product from the previous reactions in order to maximise the detection of OrM fungi. For this, the primer combination fITS9 (Ihrmark et al., 2012) and ITS4 (Tedersoo et al., 2014) were used to target the ITS2 region. These PCRs were performed with 6 μl Taq Polymerase (DreamTaq Green, Thermo Scientific, Waltham, USA), 1 μl bovine serum albumin 0.4% (w/v), 0.5 μl of 0.5 μM forward primer, 0.5 μl of 0.5 μM reverse primer, 1 μL of primary PCR product and 1 μL nuclease free water in a total volume of 10 μL. PCR conditions used were: denaturation 5 min at 94 °C, followed by 35 cycles of 30 s at 94 °C, 30 s at 55 °C and 30 s at 72 °C. The final extension time for all PCRs was 72 °C for 10 min. Each DNA extract was amplified in three replicates. All PCR products were checked on 1% agarose gel and replicates were subsequently pooled. PCR products were sent for purification and libraries analysed with paired-end sequenced using Illumina NovaSeq platform (2 x 250 bp).

### Bioinformatic analysis

The raw FASTQ files corresponding to each sequenced sample obtained from Illumina runs were processed with QIIME2 (Bolyen et al., 2019). Paired-end, demultiplexed sequence reads were imported into QIIME2 using fastq manifest format, and sequence quality control, deionizing, chimera detection and clustering into OTUs was performed using the DADA2 plugin in QIIME2 (Callahan et al., 2016). The forward and reverse sequences were truncated at 240 and 230 bp respectively, based on a minimum Phred score of 28 averaged over a 50 bp moving window. Chimeric sequences were removed performing a de novo detection using USEARCH61 (Edgar et al., 2011). After quality filtering and chimera removal, fungal operational taxonomic units (OTUs) were clustered based on 98% similarity threshold using USEARCH61. Global singletons and doubletons were removed, as they may reduce the accuracy of diversity estimates (Brown et al., 2015; Nguyen et al., 2015; Oliver et al., 2015). The closest taxonomic match of each sequence variant was then identified using the classify-sklearn function against the UNITE fungal ITS database (version 8, 16.10.2022; Abarenkov et al., 2010; Kõljalg et al., 2013). We manually checked all erroneously assigned and unidentified taxa, and we removed OTUs with a low abundance of reads from samples to avoid overestimation of diversity by using a 0.05% cut-off of the proportion of reads for each sample. Blasted sequences with a high e-value (≥ e^-50^) were considered as reliable to assign OTUs into the fungal kingdom. OTUs were regarded as OrM taxa when belonging to the families Ceratobasidiaceae, Tulasnellaceae, Sebacinaceae and Serendipitaceae with a similarity threshold of at least 85%. Subsequently, other fungal taxa such as ectomycorrhizal and saprobic fungi, known to form OrM (Dearnaley et al., 2012), were screened manually and kept for further analysis.

To infer the phylogenetic position of filtered fungal OTUs, a maximum likelihood (ML) analyses was performed in the CIPRES (Miller et al., 2011). Forty-three Tulasnellaceae sequences (OR507087 – OR507130) isolated from adult roots and protocorms of *O. italica* reported in Balducci et al. (2024) were also included for phylogenetic reconstruction. An alignment was obtained with MAFFT v.7.490 (Katoh & Toh, 2010) with default parameters and subsequently trimmed using trimAI v1.2.59 on XSEDE (Capella-Gutiérrez et al., 2009). The highest likelihood tree was calculated with RAxML-HPC2 on XSEDE and applying the GTR + GAMMA evolutionary (model v 8.2.12; Stamatakis, 2014). Branch support was calculated by nonparametric bootstrapping using 1000 pseudoreplicates. Nodes with bootstrap support ≥ 75% are considered well supported (Hillis & Bull, 1993).

### Geographical variation in OrM communities and abundance

All analyses were performed using R software version 3.5.2 (R Core Team, 2024). To test whether OrM fungi varied according to latitude and longitude, we used Generalised Additive Mixed Effect Models (GAMMs) in the ‘gamm4’ package (Wood & Scheipl, 2014). GAMMs are an extension of linear regression models that allow for analysis of non-linear effects among variables and non-normal distributions of explanatory variables. These models included a smooth term for latitude and longitude separately, while population was included as a random effect to account for variation due to sampling differences between each population. The proportion of OrM fungal OTUs detected in each population from the total number of OTUs detected were analysed using a binomial error distribution with a logit link. We found that Tulasnellaceae and Ceratobasidiaceae were widespread (see results). Therefore, we tested the influence of latitude and longitude on the proportion of (a) all OTUs in each population, (b) Tulasnellaceae OTUs only, (c) Ceratobasidiaceae OTUs only, and (d) all other OTUs per population. As six OTUs we found in every population, we used GAMMs to test whether their abundance, as measured by the number of reads per sample, also varied with latitude and longitude. For this we used a Poisson error distribution with a log link function. All model outputs from GAMMs were plotted and backtransformed from the logit or log scale.

### Multivariate analysis

Prior to the removal of non-mycorrhizal fungal OTUs, we generated rarefaction curves using the ‘rarecurve’ function in ‘vegan’ (Oksanen et al., 2019) for each population to estimate the overall coverage of the fungal communities studied (Fig. S1). Non-metric multidimensional scaling (NMDS) plots clustering each population based on OTU presence-absence were generated to visualise differences in fungal communities between regions using the Bray-Curtis coefficient as a distance measure. To test the hypotheses that mycorrhizal communities differed between regions, we used a permutational analysis of variance (PERMANOVA) with ‘adonis’ function in ‘vegan’ with 9,999 permutations, using the Bray-Curtis coefficient as a distance measure.

The effects of different environmental factors and spatial variables on OrM fungal communities were analysed by a partial Redundancy Analysis (pRDA). For this, spatial coordinates were decomposed as principle coordinates of neighbour matrices (PCNM), which generates axes that relates with variation at different spatial scales (Borcard & Legendre, 2002). The axes were produced using the ‘pcnm’ function in ‘vegan’. Four PCNM variables were produced from this analysis. Since both climatic and edaphic variables are known to be important predictors of plant species and mycorrhizal fungi assemblages at large scales (Davison et al., 2015; Tedersoo et al., 2014; Větrovský et al., 2019), we extracted regional values in Quantum GIS (QGIS Development Team, 2023) of all BIOCLIM layers from the WorldClim database (Hijmans et al., 2005) for the period 1950-2000 at 1 km^2^ resolution, and soil data from the European Soil Data Centre (ESDAC; Van Liedekerke et al., 2006) based on the Land Use and Cover Area frame Survey (LUCAS) topsoil data (Ballabio et al., 2019). To identify which soil and environmental predictors best explain the differences in OrM fungal communities, we used the ‘envfit’ function in ‘vegan’ with 9,999 permutations based on this NMDS output. Environmental parameters were log transformed to avoid bias caused by differences in unit scales. A forward selection of variables (‘forward.sel’ function in ‘vegan’) was used to select the environmental variables and spatial factors (PCNM axes and spatial coordinates), separately, that best explained variation in OrM fungal community composition. Potential collinearity between the selected predictors was examined by variance inflation factors (VIF) using ‘vif.cca’ function. Variables with VIF ≤ 5 were included. The OTU abundance matrix was Hellinger transformed (Legendre & Gallagher, 2001). To examine how OrM fungal communities varied according to key environmental predictors, we used the significant predictors from the ‘envfit’ analysis to produce a constrained ordination which included the significant spatial predictors from the PCNM analysis as conditional variables in the ordination to control for spatial differences between sampling populations. The significance of the selected variables and model was assessed by the function ‘anova.test’ in ‘vegan’.

### Hierarchical Modeling of Species Communities

As OrM fungi frequently form communities that may respond similarly to environmental variables, we tested whether their co-occurrences can be explained by abiotic variables or whether there were residual co-occurrences among OrM fungi that were unexplained by abiotic variables. There are several frameworks to infer how species jointly respond to environmental variables using Joint Species Distribution Modelling (JSDM) (Pollock et al., 2014; Warton et al., 2015). These approaches are based on multivariate regression and allow partitioning of shared environmental responses among species from residual correlations among species. The Hierarchical Modelling of Species Communities (HMSC) framework is a joint species distribution model that allows a set of covariates to be modelled on multiple response variables (Ovaskainen et al., 2017). This framework is a hierarchical generalized linear mixed model and is structured by both fixed effects and random effects, allowing to jointly model the occurrences of OrM OTUs identified in each individual *O. italica* as a function of environmental covariates (Tikhonov et al., 2017). Hence, we can estimate to what extent co-occurrence of OrM OTUs is explained by environmental variables and interactions between OTUs for which statistically supported co-occurrence remains after accounting for environmental variables.

### JSDM model fitting

We screened OTUs that occurred in at least 10% of *O. italica* roots, resulting in 38 OTUs belonging to six OrM fungal families (Ceratobasidiaceae, Inocybaceae, Russulaceae, Sebacinaceae, Serendipitaceae, and Thelephoraceae) were retained for the HMSC analysis. This is because replication of OTUs among orchid individuals is necessary for HMSC to quantify co-occurrence among OrM taxa (Ovaskainen & Abrego, 2020; Abrego et al., 2020). We fitted two models; a “raw co-occurrence” model where sequencing depth (the log-transformed number of sequences) was used as the only explanatory variable as a ‘null model’, and a “residual co-occurrence” model. Variables included; (i) organic phosphorus, (ii) annual mean temperature (bio1), (iii) mean precipitation in the wettest quarter (bio16), (iv) mean precipitation in the driest quarter (bio17), (v) PCNM1, (vi) PCNM2 (two spatial variables to control for spatial correlation), (vii) elevation, and (viii) elevation2 (to control for nonlinear responses of fungal communities to elevation). To account for sampling effects, we also included sequencing depth (as in the raw co-occurrence model) as an explanatory variable. To account for zero-inflation in the data, we used a hurdle model approach consisting of two parts; (i) the presence-absence of OTUs using a probit regression and, (ii) the relative abundance of OrM taxa, measured as the number of sequences per OTU, a lognormal model conditional on presence (i.e., where absences were removed) (Ovaskainen & Abrego, 2020). Therefore, four outputs emerge from the hurdle approaches: the (i) “raw probit model”, (ii) the “residual probit model”, (iii) the “raw lognormal model”, and (iv) the “residual lognormal model”. We used the package ‘Hmsc’ (Tikhonov et al., 2020) assuming the default prior distributions (Ovaskainen & Abrego, 2020).

We fitted our models with three chains of 100,000 Markov chain Monte Carlo (MCMC) iterations (a total of 300,000 MCMC iterations) comprised of 1,000 MCMC samples, with the samples thinned by a factor of 100, out of which we discarded the first 50,000 samples as burn-in. We assessed MCMC convergence by examining the potential scale reduction factors (Gelman & Rubin, 1992) of the model parameters. We examined residual co-occurrence between pairs of OrM OTUs according to a 0.9 posterior support of the association for all four HMSC models. Based on the outputs of the residual probit and residual lognormal model, we calculated the proportion of variance explained by each environmental variable for each model using the variance partitioning (VP) function in ‘Hmsc’.

## Results

We retrieved 340 putative OrM fungal OTUs after quality filtering and removal of aberrant, unidentifiable low-quality sequences. Rarefaction curves revealed sufficient sampling effort for the detection of total fungal diversity for each population (Fig. S1). After applying a 0.05% abundance threshold, our final dataset consisted of 80 OrM fungal OTUs (2,371,010 sequences) from seven mycorrhizal families associated with *O. italica* populations. Phylogenetic analysis revealed that the OrM fungal OTUs detected in this study are dispersed among multiple clades, including those of known orchid mycobionts (Fig. S2 – S7). Representative sequences for each mycorrhizal OTU found in this study were submitted to GenBank nucleotide database under accession numbers PP935024 - PP935103.

The OrM fungal OTUs were primarily dominated by the rhizoctonia-like fungi assigned to the following families: Tulasnellaceae (31 OTUs; 2,094,142 sequences), Ceratobasidiaceae (15 OTUs; 236,060 sequences) and Serendipitaceae (six OTUs, 8,278 sequences). The remaining OrM fungal OTUs, albeit at much lower abundances, belonged to four families with putative mycorrhizal abilities: Sebacinaceae (eight OTUs, 5,423 sequences), Thelephoraceae (10 OTUs, 21,278 sequences), Russulaceae (six OTUs, 3,706 sequences) and Inocybaceae (four OTUs, 2,123 sequences). 88.32% of reads belonged to Tulasnellaceae indicating that they were the most abundant mycorrhizal family associated with *O. italica*. Six OTUs were recovered from all 23 populations, five of which belonged to Tulasnellaceae (OTUs tul26, tul27, tul28, tul29, and tul30) and one from Ceratobasidiaceae (cer15).

The proportion of OrM OTUs per population increased with increasing longitude (ꭓ^2^ = 18.06; p < 0.001; Fig. 2a) but did not vary according to latitude (ꭓ^2^ = 0.989; p = 0.32; Fig. S8a). Likewise, the proportion of Tulasnellaceae OTUs (ꭓ^2^ = 7.423; p < 0.001; Fig. 2b) and Ceratobasidiaceae OTUs (ꭓ^2^ = 11.45; p < 0.001; Fig. 2c) increased with increasing longitude but did not vary with latitude (ꭓ^2^ = 0.359; p = 0.549 and ꭓ^2^ = 0.523; p = 0.47 respectively; Fig. S8b & c). The strength of both the longitude (Fig. S9) and latitude (Fig. S10) gradients did not change when we removed the two Greek populations. The number of reads per sample of tul26 (ꭓ^2^ = 4.544; p = 0.033), tul27 (ꭓ^2^ = 30.93; p < 0.001), tul28 (ꭓ^2^ = 26.38; p < 0.001), tul29 (ꭓ^2^ = 19.14; p < 0.001), tul30 (ꭓ^2^ = 20.97; p < 0.001) and cer15 (ꭓ^2^ = 4.172; p = 0.041) increased with increasing longitude (Fig. 3). The number of reads per sample for tul26 (ꭓ^2^ = 14.27; p < 0.001), tul27 (ꭓ^2^ = 4.189; p = 0.041), tul29 (ꭓ^2^ = 5.276; p = 0.022), tul30 (ꭓ^2^ = 7.025; p = 0.008) and cer15 (ꭓ^2^ = 5.042; p = 0.025) decreased with increasing latitude, while the abundance of tul28 (ꭓ^2^ = 3.384; p = 0.066) marginally did not vary according to latitude (Fig. 4). After removing the two Greek populations, the number of reads per sample for tul28 (ꭓ^2^ = 5.544; p = 0.019), tul29 (ꭓ^2^ = 3.682; p = 0.050) and tul30 (ꭓ^2^ = 3.858; p = 0.049) increased with increasing longitude (Fig. S11) while both tul26 (ꭓ^2^ = 10.38; p = 0.001) and tul30 (ꭓ^2^ = 4.307; p = 0.038) decreased with increasing latitude (Fig. S12).

**Fig. 2.**
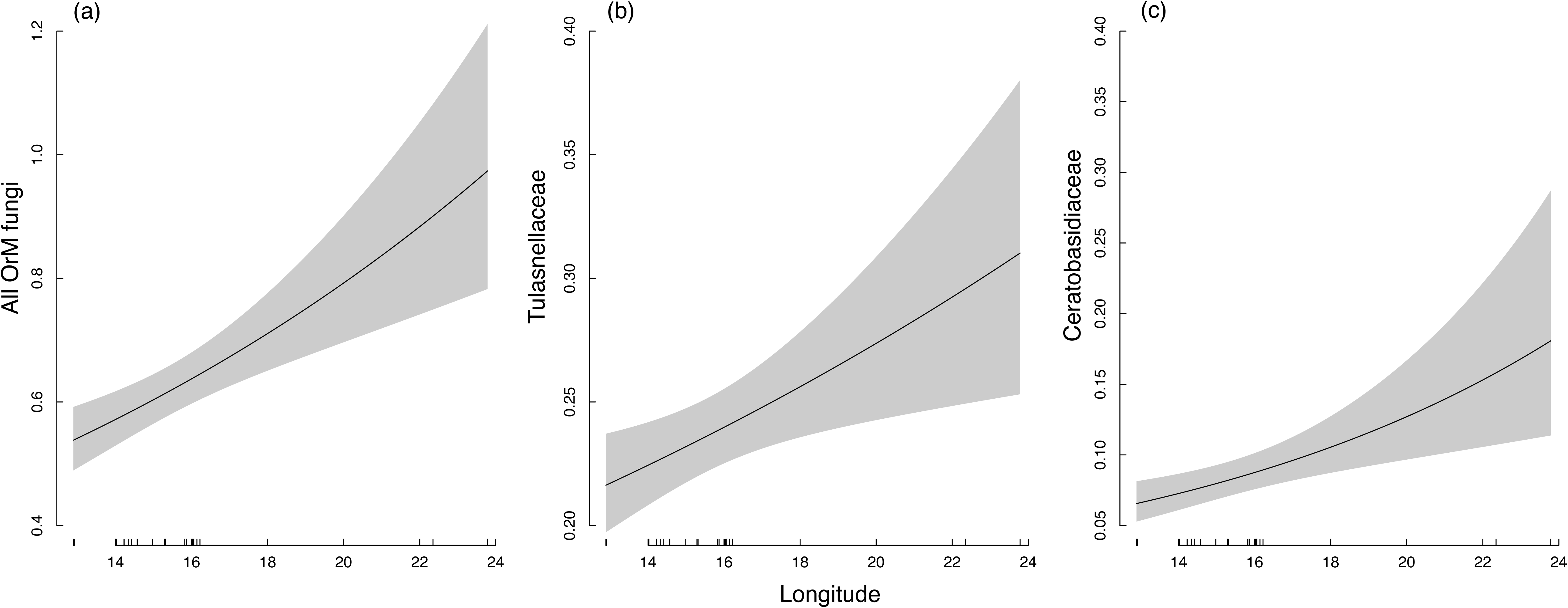
The influence of longitude on the proportion of OrM OTUs in each population for; (a) all OTUs, (b) Tulasnellaceae OTUs and (c) Ceratobasidiaceae OTUs. Model outputs are back-transformed from the logit scale.

**Fig. 3.**
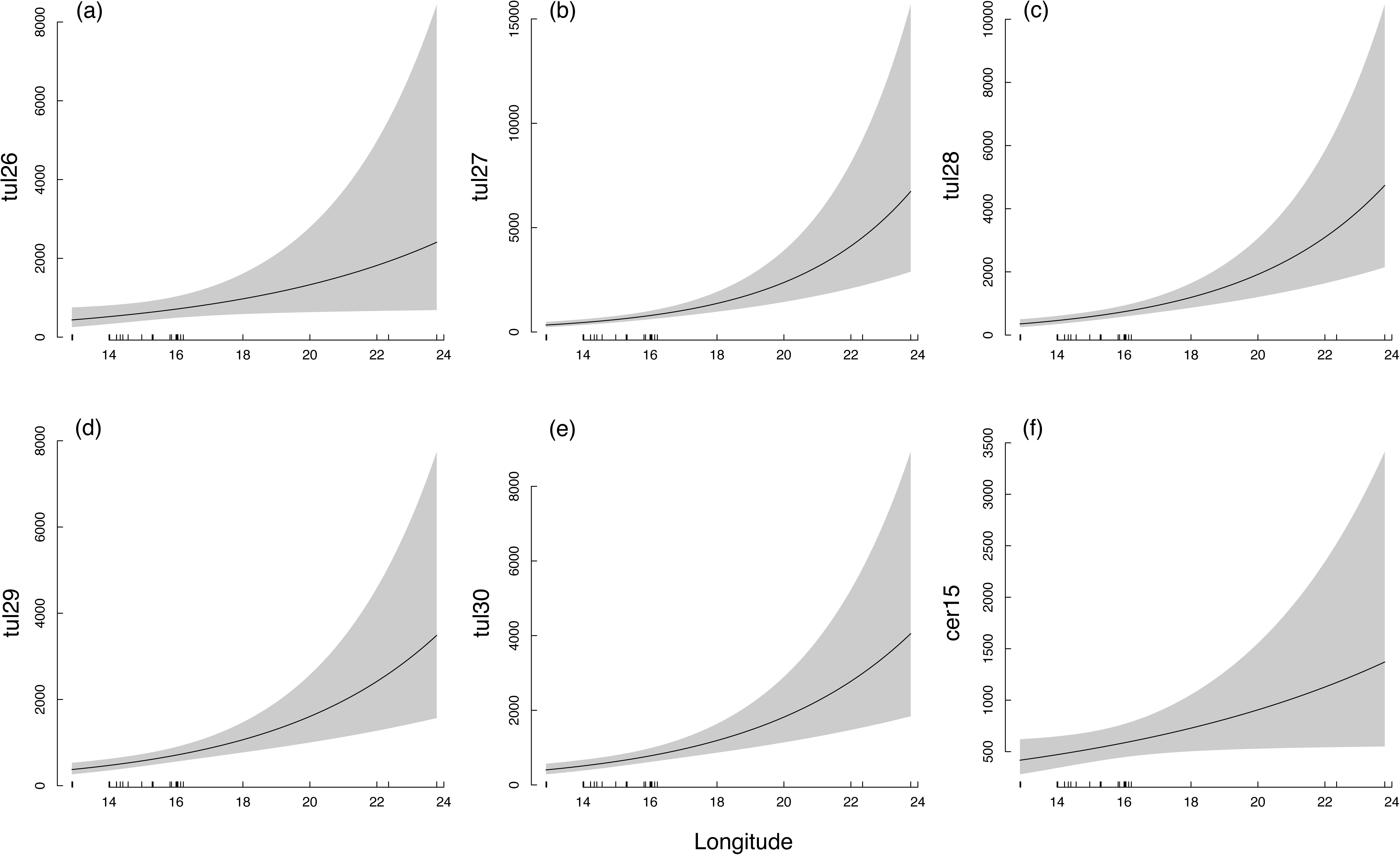
Influence of longitude on the mean number of reads of; (a) tul26, (b) tul27, (c) tul28, (d) tul29, (e) tul30 and (f) cer15. Model outputs are back-transformed from the log scale.

**Fig. 4.**
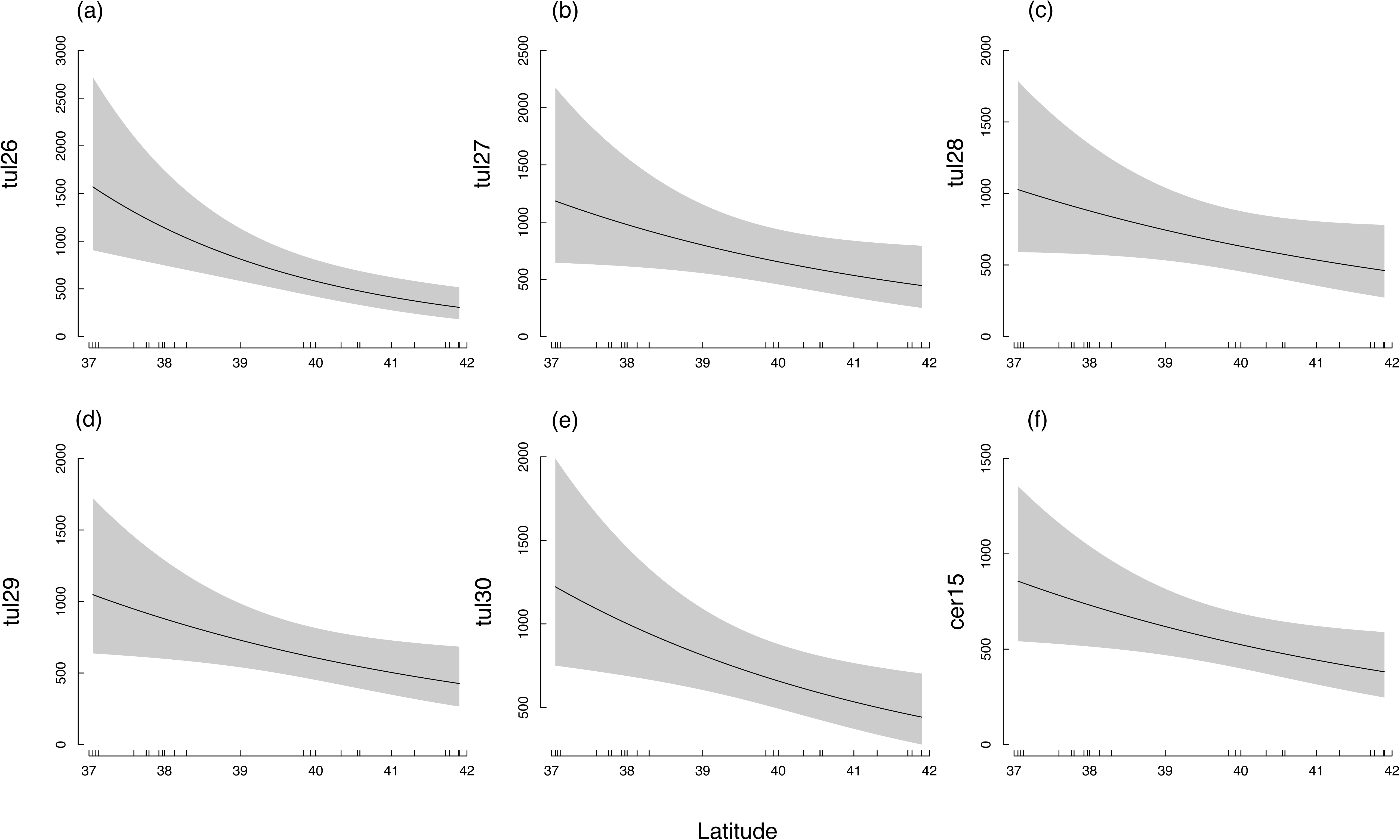
The influence of latitude on the mean number of reads of; (a) tul26, (b) tul27, (c) tul28, (d) tul29, (e) tul30 and (f) cer15. Model outputs are back-transformed from the log scale.

The NMDS showed that mycorrhizal community composition typically clustered together according to region (Fig. 5). PERMANOVA supported the NMDS and revealed that mycorrhizal communities differed between regions (R^2^ =0.673, F = 9.27, p < 0.001). After controlling for spatial variability, the partial RDA showed that mycorrhizal communities varied according to different environmental variables (RDA model: adjusted *R*^2^ = 0.33; p = 0.001; Fig. 6). These were; (i) annual mean temperature, (ii) mean precipitation in the driest quarter, (iii) mean precipitation in the wettest quarter, (iv) organic phosphorous and (v) two PCNM variables (PCNM 1 and PCNM 2). OrM fungal communities in populations from Puglia differed from other regions according to soil phosphorous (p = 0.024; Fig. 6). Populations from Greece and Sicily differed according to annual mean temperature (p = 0.001), while both precipitation in the driest quarter (p = 0.002) and precipitation in the wettest quarter (p = 0.002) were significant but displayed a more varied pattern in ordination space among the remaining regions (Fig. 6).

**Fig. 5.**
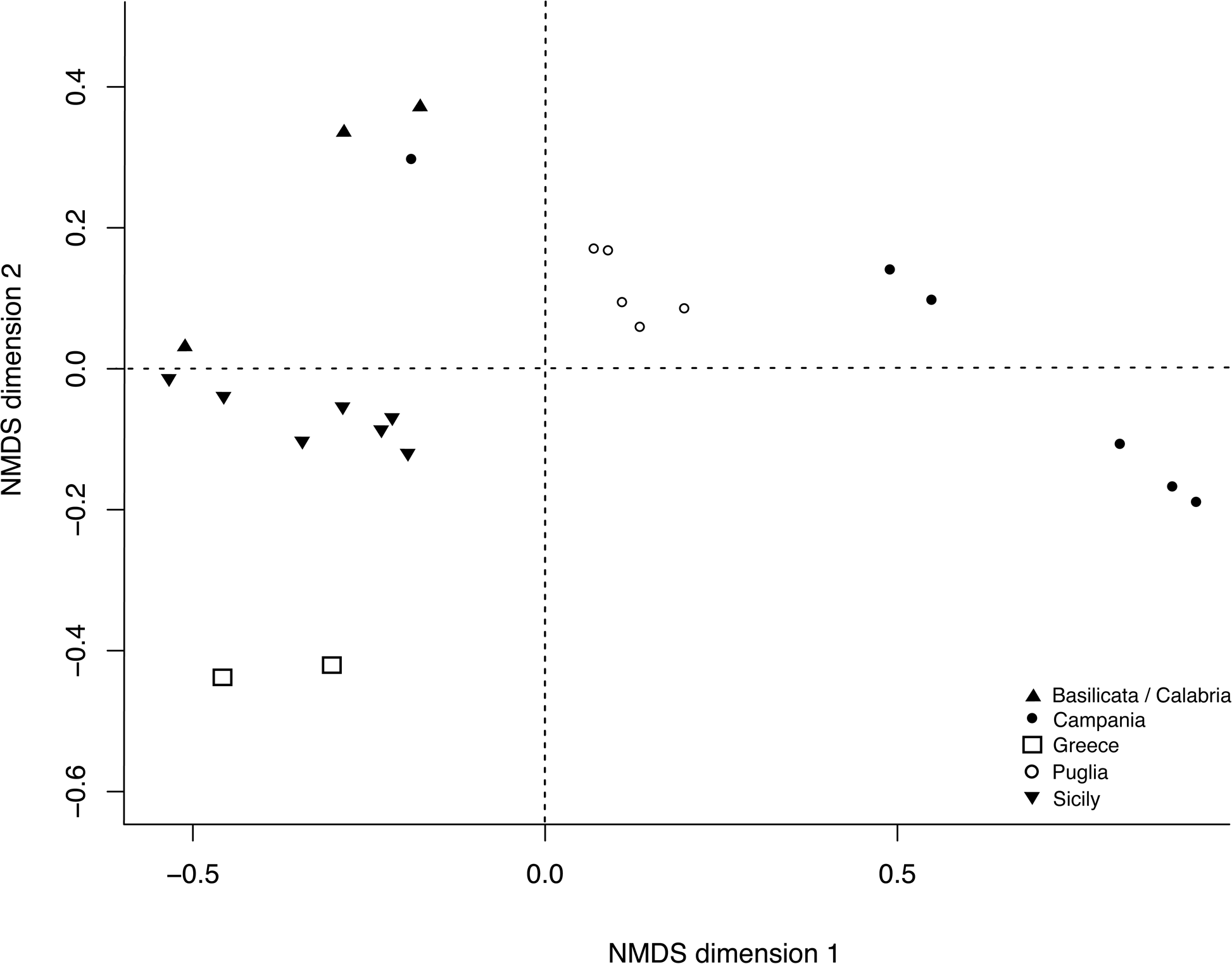
NMDS ordination plot showing the differences between *Orchis italica* populations in their orchid mycorrhizal fungi composition. Stress = 0.077.

**Fig. 6.**
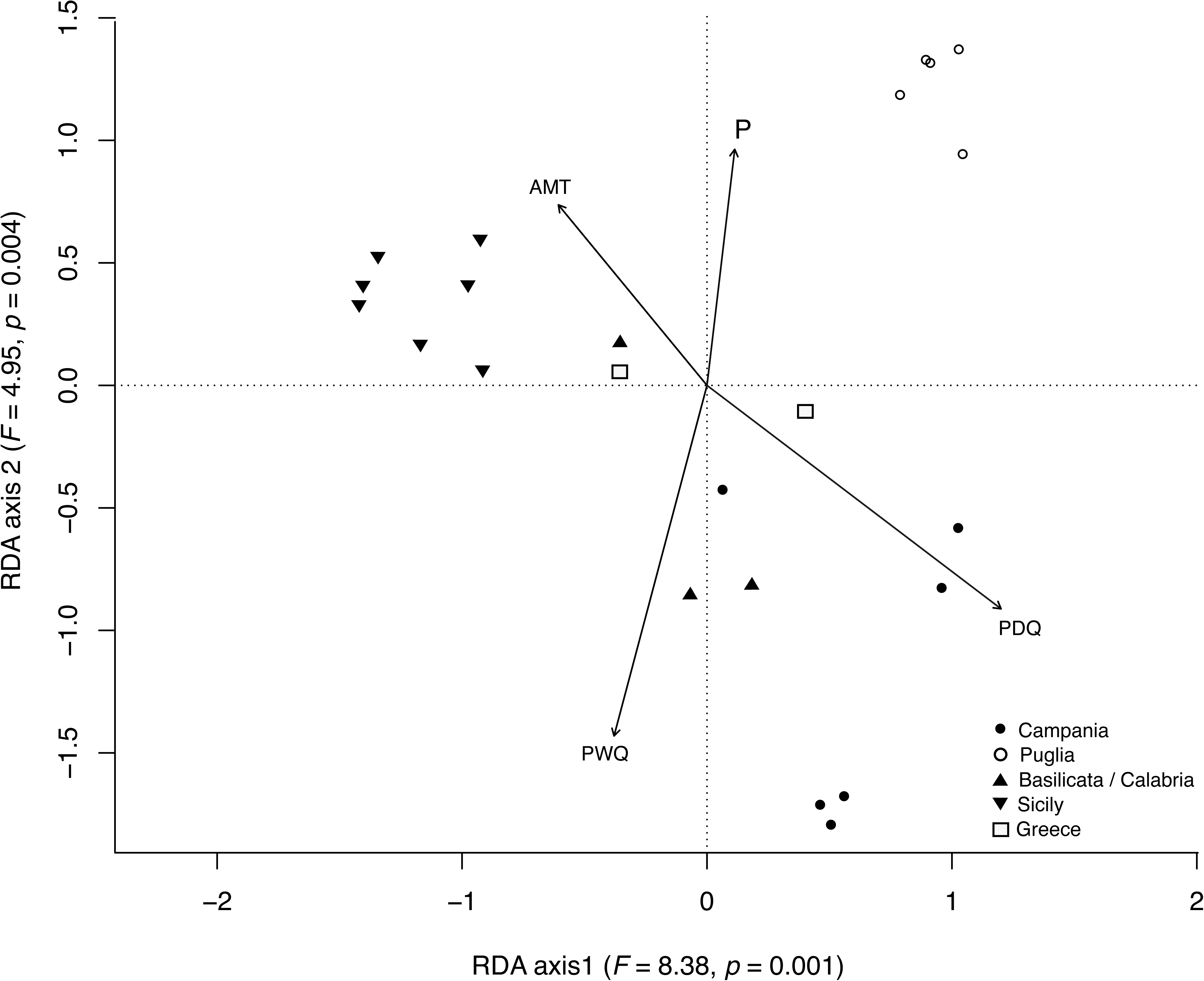
Partial Redundancy analysis (RDA) showing the effects of annual mean temperature (AMT), precipitation in the driest quarter (PDQ), precipitation in the wettest quarter (PWQ) and phosphorous on orchid mycorrhizal fungal community between populations of *Orchis italica*. The length of the vectors represents the strength of that variable on the distribution of fungal taxa.

All HMSC models performed well, with potential scale reduction factor for each model approximately 1, and visual inspection of posterior trace plots showing the MCMC chains yield nearly identical results and mixed well. Variance partitioning of the presence-absence probit model revealed that the spatial variable PCNM1, annual mean temperature, and precipitation in the driest quarter were the most important variables in explaining co-occurrence based on presence-absence data (Table 1). However, variance partitioning of the lognormal conditional on presence model revealed that spatial variables were not as important, but that sampling effort (i.e., number of sequences) and annual mean temperature were the most important variables in determining OrM fungal co-occurrences (Table 1). Both the probit and lognormal conditional-on-presence models that included only number of reads as an explanatory variable had relatively high numbers of positive and negative residual co-occurrences by chance (Fig. 7a & b). Residual models that included environmental variables of both the presence-absence and lognormal conditional on presence datasets containing environmental variables revealed fewer residual co-occurrences (Fig 7c & d). In the residual probit model the co-occurrence patterns based on presence-absence data showed that most Tulasnellaceae OTUs co-occurred frequently, while the key six OTUs did not co-occur. However, in the residual lognormal conditional on presence model, there were positive and negative co-occurrences among these ubiquitous OTUs (Fig. 7).

**Fig. 7.**
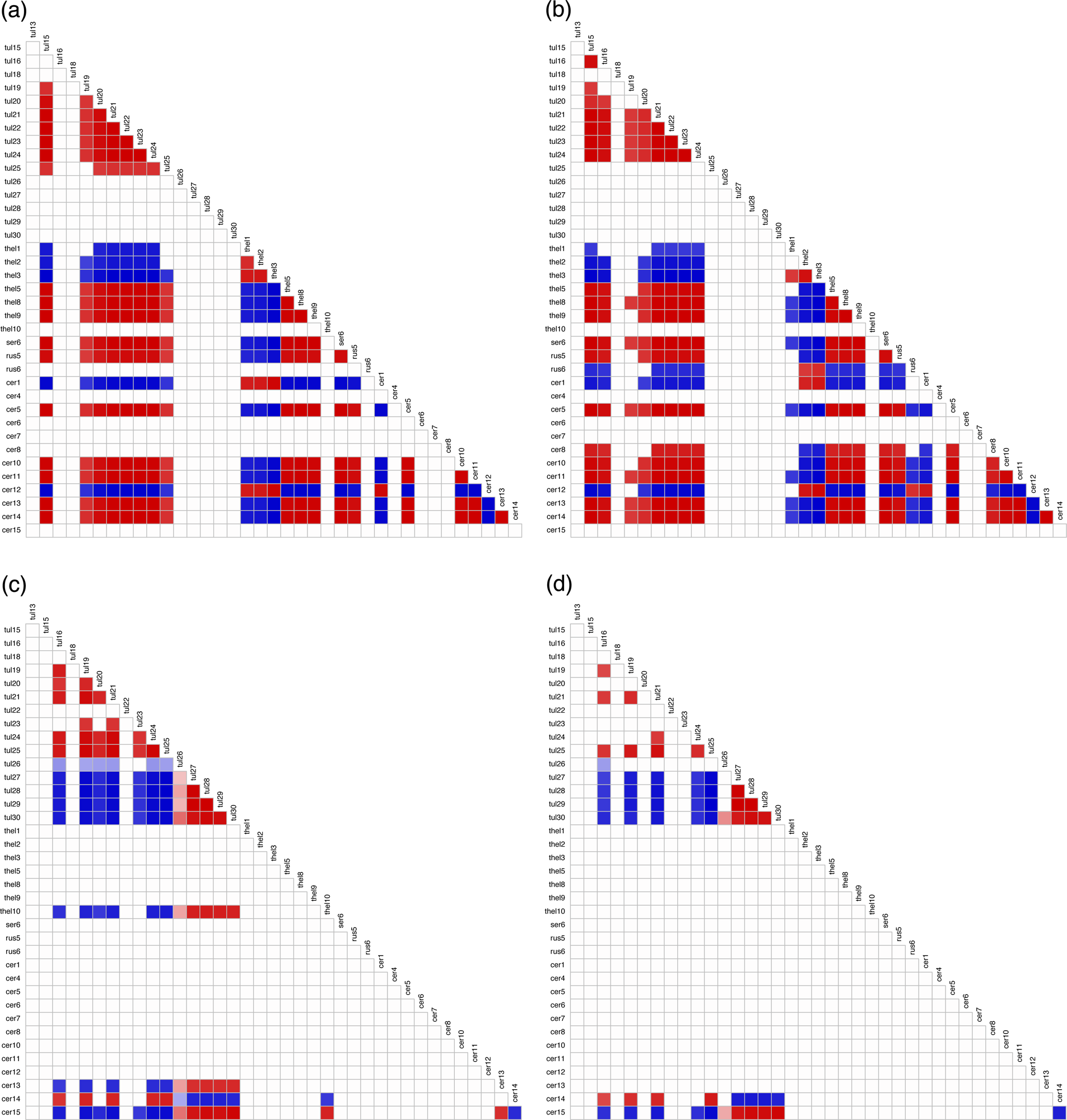
Co-occurrence patterns among OrM fungi associated with *Orchis italica* based on (a) the raw probit model based on number of reads only, (b) the residual probit model, (c) the raw lognormal model based on number of reads only, and (d) the residual lognormal model, with 0.9 posterior statistical support. Red squares represent positive residual correlations, blue squares represent negative residual correlation.

**Table 1.** Variance partitioning of the percent contribution of fixed and random effects in the residual co-occurrence probit model and the residual co-occurrence conditional on presence lognormal model.

|  | Variable | Probit model (%) | Lognormal model (%) |
| --- | --- | --- | --- |
| Random effect | Plant | 12.0 | 10.1 |
| Fixed effects | Elevation | 10.0 | 6.1 |
|  | Elevation<sup>2</sup> | 9.4 | 9.7 |
|  | PCNM1 | 14.2 | 6.4 |
|  | PCNM2 | 3.8 | 4.5 |
|  | Annual mean temperature (bio1) | 13.8 | 11.6 |
|  | Precipitation driest quarter (bio16) | 14.2 | 4.2 |
|  | Precipitation wettest quarter (bio17) | 6.7 | 5.0 |
|  | Phosphorous | 2.7 | 3.6 |
|  | Number of reads | 13.3 | 38.7 |

## Discussion

The results of this study show that geographical distance influences OrM fungal community composition associated with the Mediterranean orchid, *O*. *italica*. The diversity and composition of OrM fungal communities differed among regions and were correlated with variation in both climatic (mean annual temperature, precipitation in the driest quarter and precipitation in the wettest quarter) and soil variables. This is consistent with previous studies that have shown that mean annual temperature and precipitation affect both fungal diversity and soil community composition in the soil (Rasmussen et al., 2018; Tedersoo et al., 2012; Toljander et al., 2006). High temperatures have been shown to increase the activity of saprotrophic fungi, enabling them to obtain more carbon to support the growth of orchids (Shefferson et al., 2019). Furthermore, some dominant OrM fungi in roots of *O. italica* such as members of the Tulasnellaceae are positively affected by increased rainfall (Jasinge et al., 2018). Soil chemical variables may affect OrM fungal communities (Mujica et al., 2016; Vogt-Schilb et al., 2020). We found that OrM fungal communities differ substantially according to phosphorous, particularly in populations from the Puglia region. This could be explained by an increase in soil phosphorous concentration and the number of Ceratobasidiaceae OTUs detected here compared to other regions. Similarly, Mujica *et al*. (2016) found a higher abundance of Ceratobasidiaceae OTUs (instead of Tulasnellaceae OTUs) in the roots of *Bipinnula fimbriata* at locations with high phosphorus concentration. Hence, OrM fungal communities may be the result of complex interactions between extrinsic factors such as geographical location, climatic and edaphic conditions.

Our data support previous studies showing that members belonging to Tulasnellaceae are the primary fungal associates in *Orchis* (Girlanda et al., 2011; Jacquemyn et al., 2010, 2011; Mennicken et al., 2023). We found an increase in OrM fungal community diversity with increasing longitude. Longitudinal variation in species distributions is poorly studied, however in the Mediterranean, east-west variation should be expected to be an important factor determining species occurrences. This is because the western part of the Mediterranean may have undergone a post-glacial recolonization from eastern refugia during the past 20,000 years (Médail & Diadema, 2009). There have also been clines in climate severity during the last inter-glacial cycle in the Mediterranean (Conord et al., 2012; Fady & Conord, 2010). In contrast, patterns in biodiversity are primarily described in terms of latitude, with the expectation that there is a reduction in abundance, diversity, or a combination of both with increasing distance from the Equator (Hillebrand, 2004; Kinlock et al., 2018). We found no evidence between OrM fungal diversity and latitude. These results contrast Duffy *et al*. (2019) who showed that OrM fungal diversity decreases with increasing latitude in Europe. However, this result may be because we sampled over a smaller latitude range compared with Duffy *et al*. (2019) and it could be that reductions in diversity with latitude exist over larger latitudinal gradients.

Five of the 30 Tulasnellaceae OTUs and one of the 15 Ceratobasidiaceae OTUs were detected in all 23 populations, revealing that these OrM fungi have broad distributions and may tolerate a wide range of climatic and edaphic conditions. The abundance, as measured by the number of reads, of these key OrM fungal taxa decreased with increasing latitude and increased with increasing longitude. As one of these OTUs (tul25) may be important in the germination of *O. italica* (Balducci et al., 2024), it could be that the reduction in the abundance of these OTUs with latitude reflects the limit of *O. italica* in the Mediterranean. *Orchis italica* does not occur in northern Europe and it may be that these OrM fungi are absent or occur at such low abundance that it renders it impossible for *O. italica* to establish outside of its current range. Likewise, as these fungal taxa increase with increasing longitude (i.e., towards the east), it could be that populations should be more abundant in the eastern part of the range of *O. italica*. It will be necessary to experimentally translocate *O. italica* outside of its native range to fully understand the limits to its distribution.

Instead, OrM fungal occurrence and communities may ultimately be determined by both abiotic (i.e., climatic and soil characteristics) and biotic factors such as competition and facilitation. We found that the presence-absence co-occurrence of OTUs was determined mainly by spatial factors (PCNM1), annual mean temperature, and precipitation in the driest quarter. Likewise, the abundance of OTUs when present was influenced by annual mean temperature. The residual co-occurrences from the presence-absence probit HMSC model indicate the presence of positive co-occurrences among Tulasnellaceae OTUs, and positive and negative co-occurrences among Ceratobasidiaceae, Thelephoraceae, Russulaceae OTUs. These co-occurrences may represent casual associations among OTUs when they occur in different populations, while the six common OTUs did not have any residual co-occurrences, likely reflecting their ubiquitous occurrence throughout the range of *O. italica*. However, the lognormal conditional on presence HMSC model revealed that these ubiquitous Tulasnellaceae OTUs may facilitate their co-occurrence or compete both with each other and Ceratobasidiaceae OTUs. These positive and negative interactions between OTUs can be expected to have played a major role in the assembly of root associated fungal communities within adult roots. Nevertheless, this is interpreted with caution as it is somewhat controversial to infer ecological mechanisms using corelative statistical approaches such as residual co-occurrences among OTUs (Blanchet et al., 2020).

Shefferson *et al*. (2019) suggested that interactions between orchids and mycorrhizal fungi may represent a case of apparent generalism, whereby orchids frequently associate with many mycorrhizal taxa, but only a subset of other taxa contribute unique or substantial resources to the orchid. In line with the results of Calevo and Duffy (2023) it could be that there is a temporal succession of OrM fungi colonization of orchid roots. It is not clear whether OrM taxa other than Tulasnellaceae play equally important roles in the germination and subsequent establishment of *O. italica* as a fully photosynthetic individual (but see Balducci et al., 2024). Such partitioning of space on the orchid roots may result in both positive and negative residual co-occurrences detected here. Similar mechanisms may be present in other orchids associating with a core set of OrM fungi (Bidartondo et al., 2004; Esposito et al., 2016). However, in other orchid species (e.g., *Epipactis*), there is almost complete turnover in mycorrhizal taxa across large spatial scales (Xing et al., 2020). Ventre Lespiaucq *et al*. (2021) proposed that OrM associations with an orchid could be explained by fidelity to a narrow range of OrM but with noise, functional turnover of OrM, and environmentally driven turnover. These mechanisms could lead to the existence of a core set of fungi associating with most individuals of an orchid species. In our case, partial OrM turnover may be environmentally driven, but fidelity with noise is likely as *O. italica* has a narrow specialization on a core group of fungi that may be involved in key phases of its life cycle. We propose that future studies quantify how key OrM fungi co-occur spatially and temporally, and perform experiments to understand the role of abiotic variables in the growth of fungi (Jacquemyn et al., 2017; van Der Heijden et al., 2015).

## Conclusions

In conclusion, this study has shown that both spatial and environmental factors were important predictors in structuring OrM fungal communities across large geographical areas. This highlights the importance of quantifying and identifying the distribution of these mycorrhizal associates as well as their response to climatic and edaphic factors in understanding the current and future distribution of *O. italica*.

## Acknowledgements

We would like to thank Spyridon Oikonomidis for contributing samples from Greece for the molecular analyses and sequencing. We gratefully acknowledge funding from the Italian Ministry of Research (PRIN grant number 2017TP3SJL).

## Data availability statement

We confirm that, should the manuscript be accepted, the data supporting the results will be archived in Dryad and the data DOI will be included at the end of the article.

## Supporting information

Additional supporting information may be found online in the Supporting Information section at the end of the article.

