## Supplementary figures and images for "Orchid mycorrhizal communities associated with *Orchis italica* are shaped by ecological factors and geographical gradients"

### Fig. S1

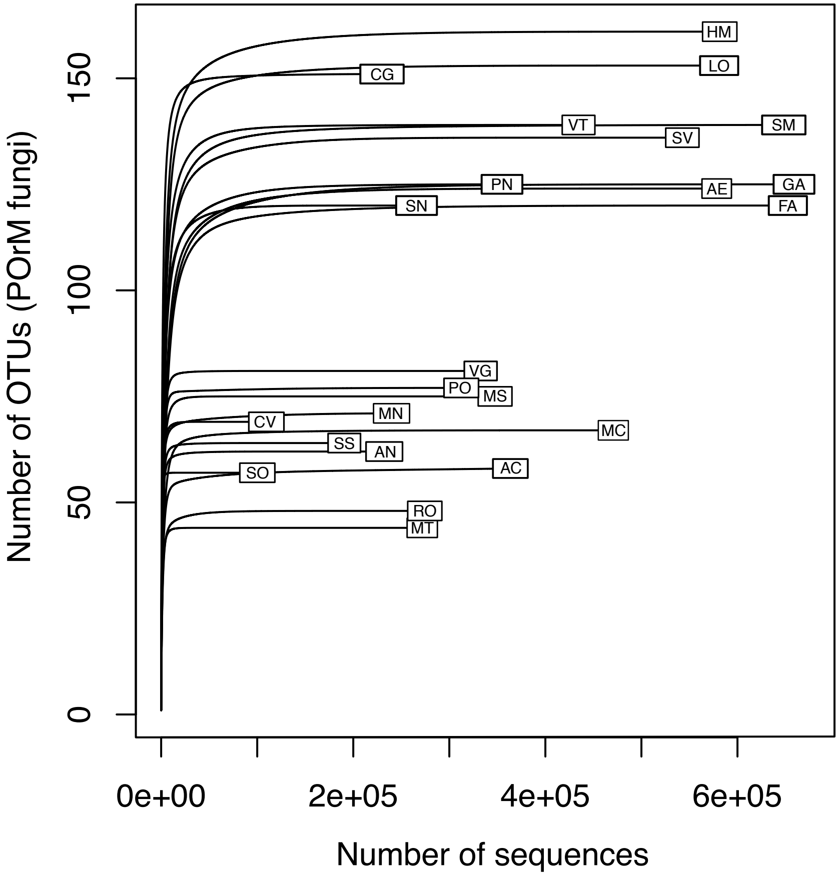

### Fig. S2

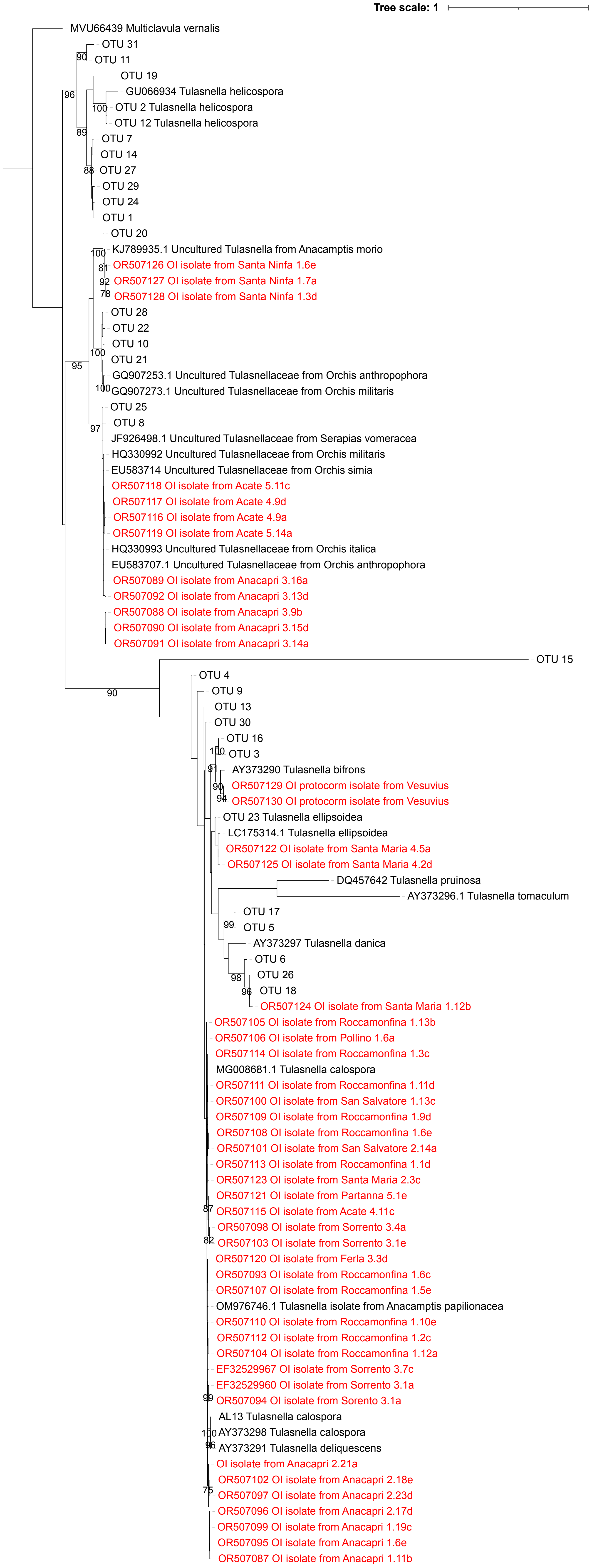

### Fig. S3

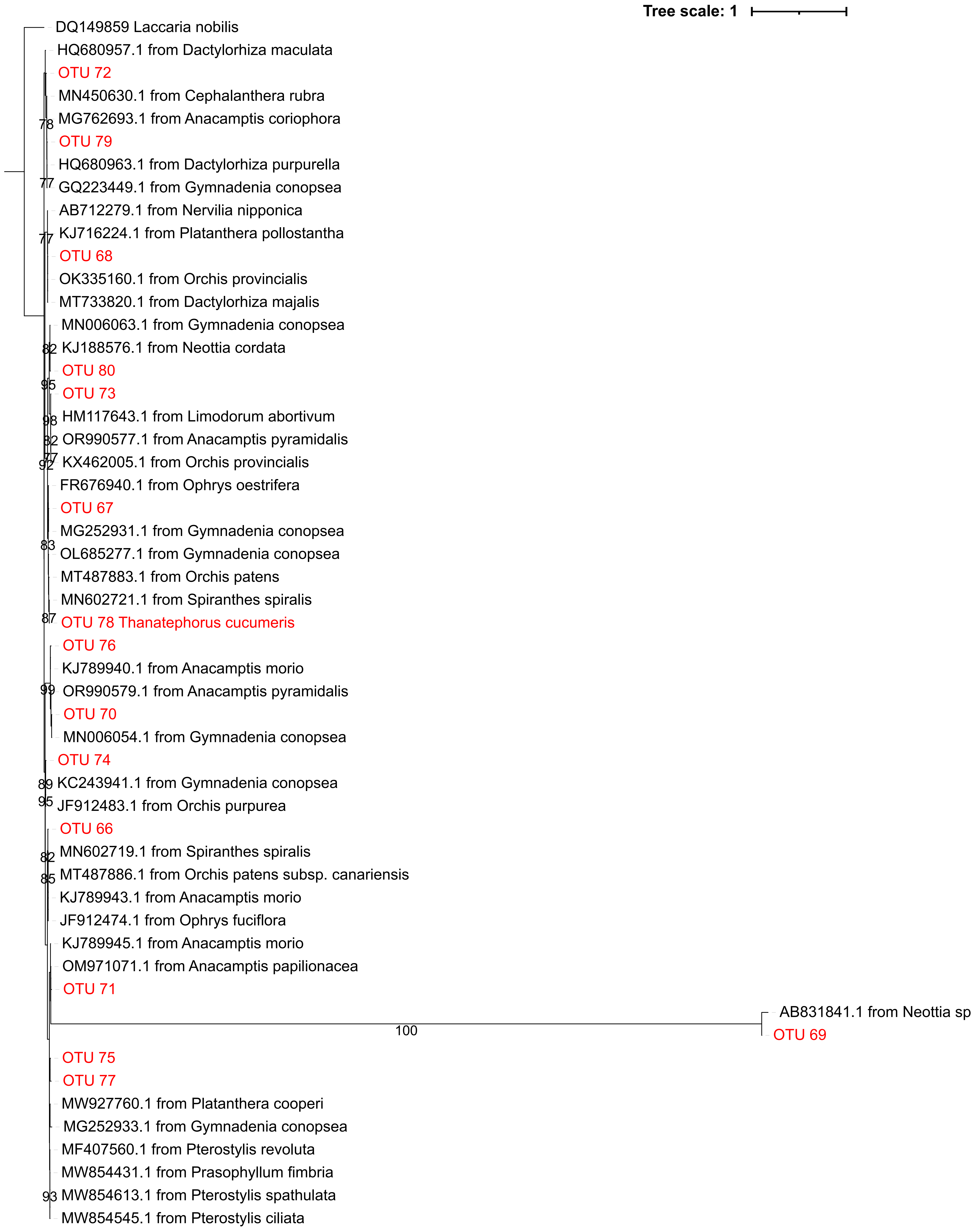

### Fig. S4

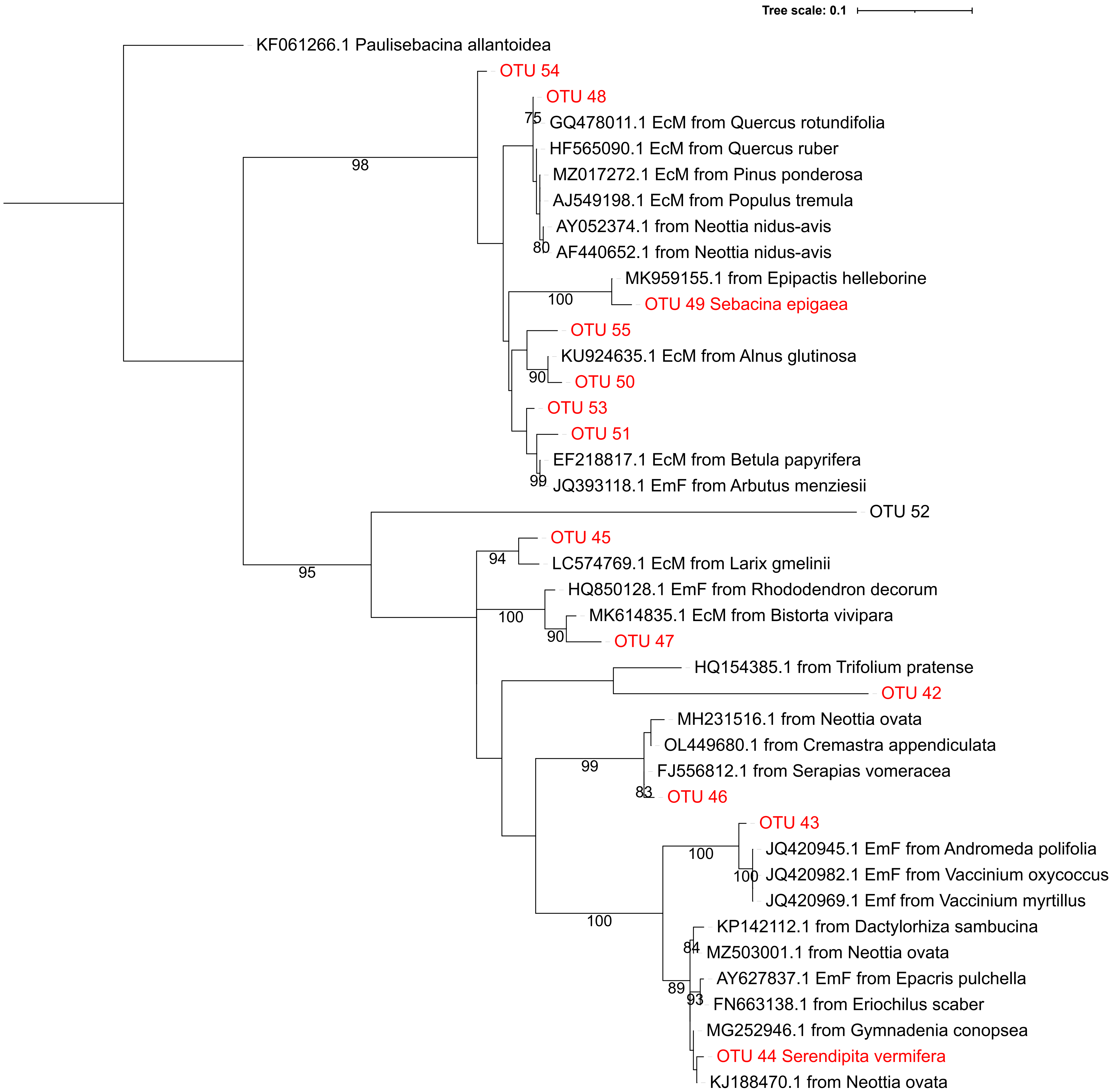

### Fig. S5

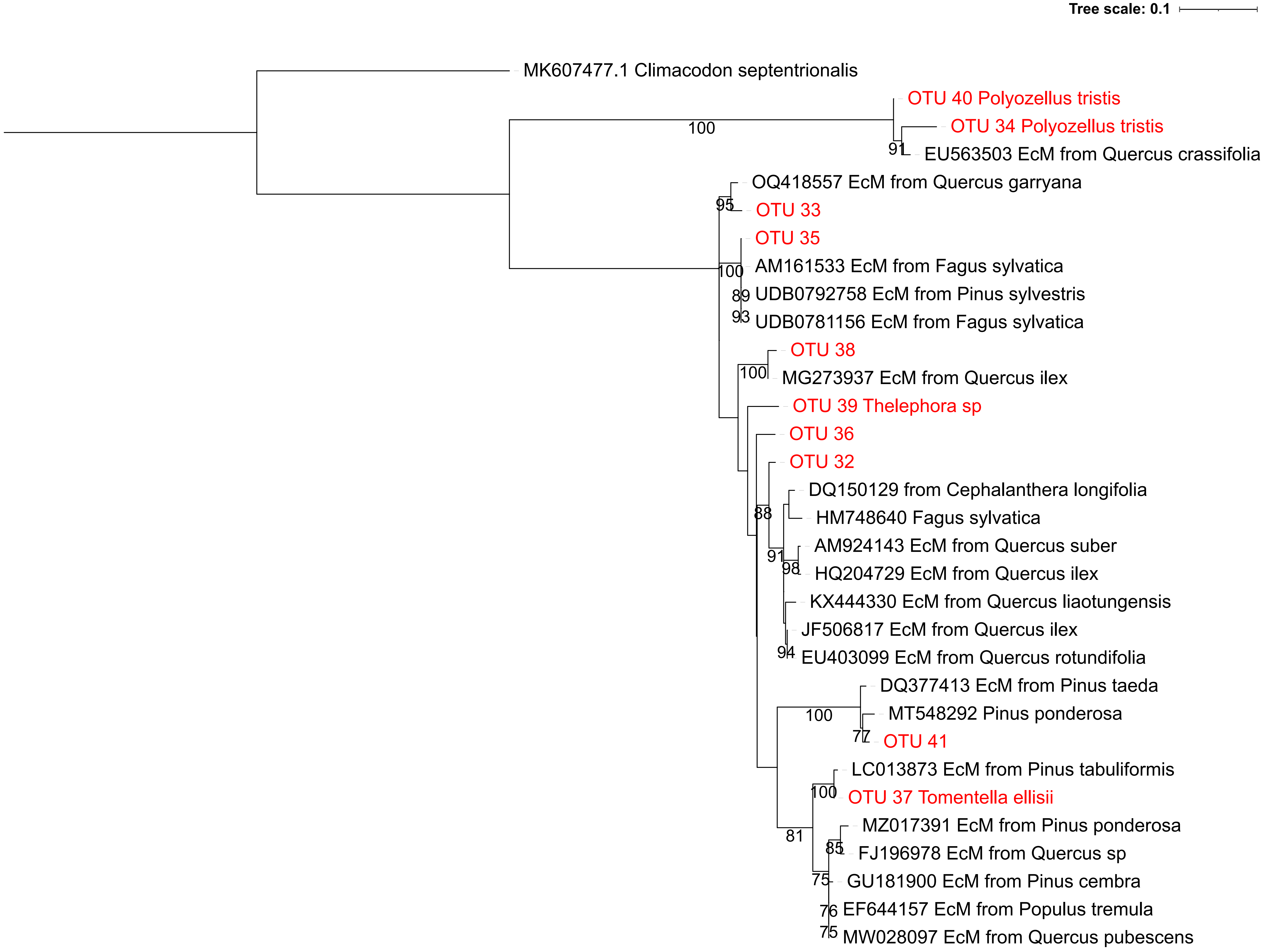

### Fig. S6

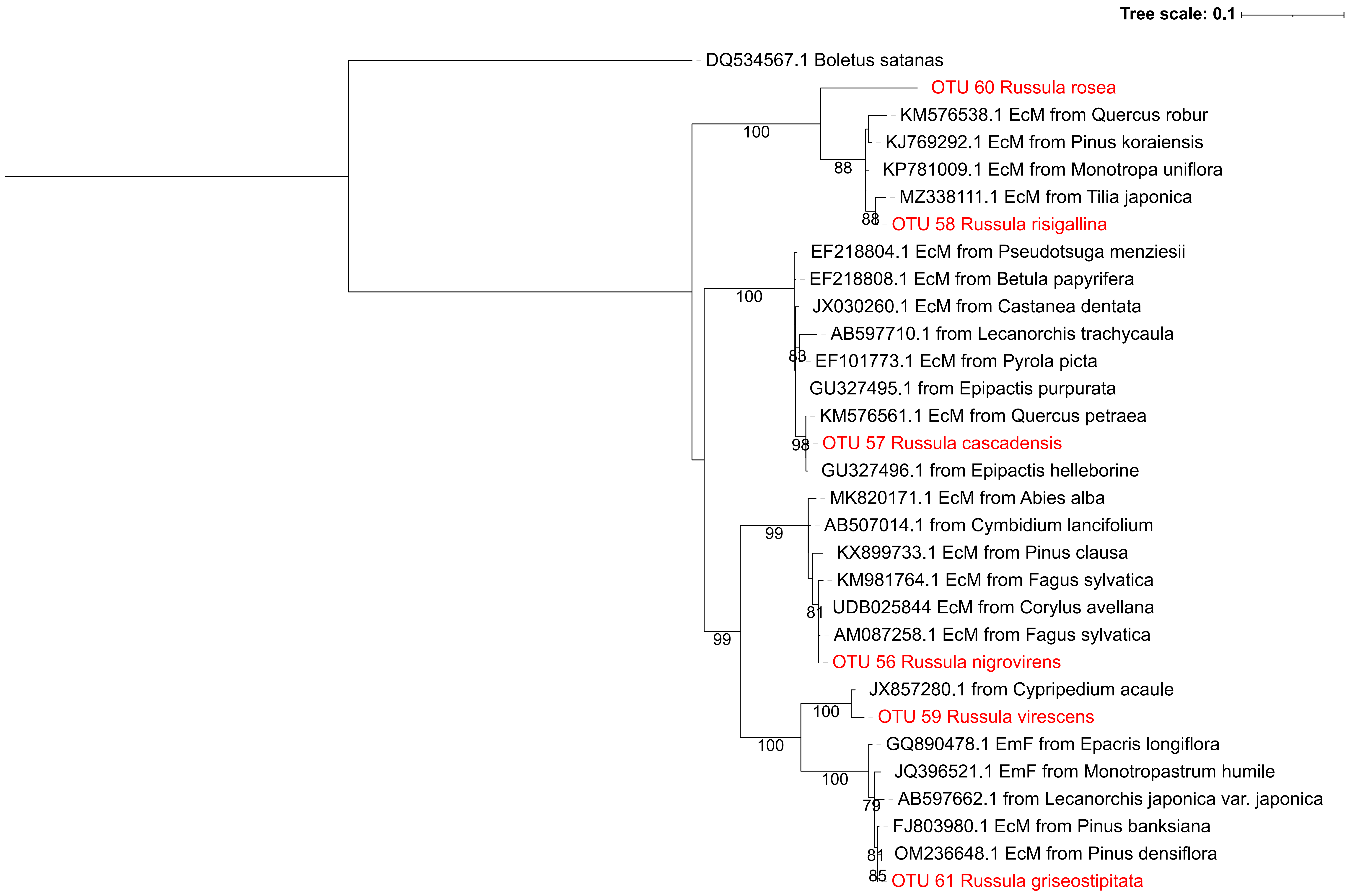

### Fig. S7

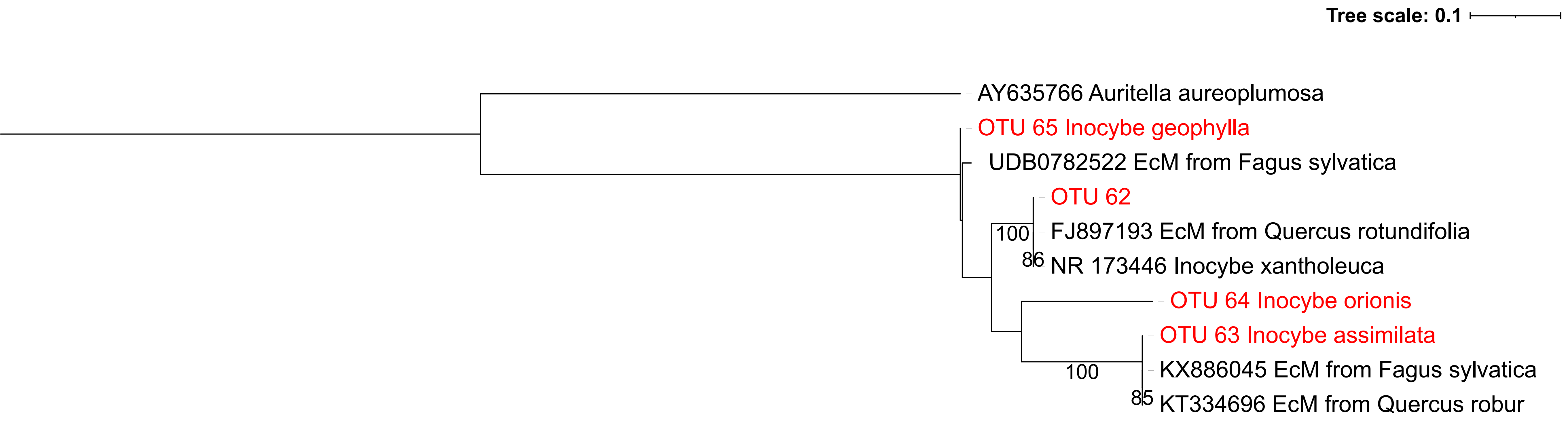

### Fig. S8

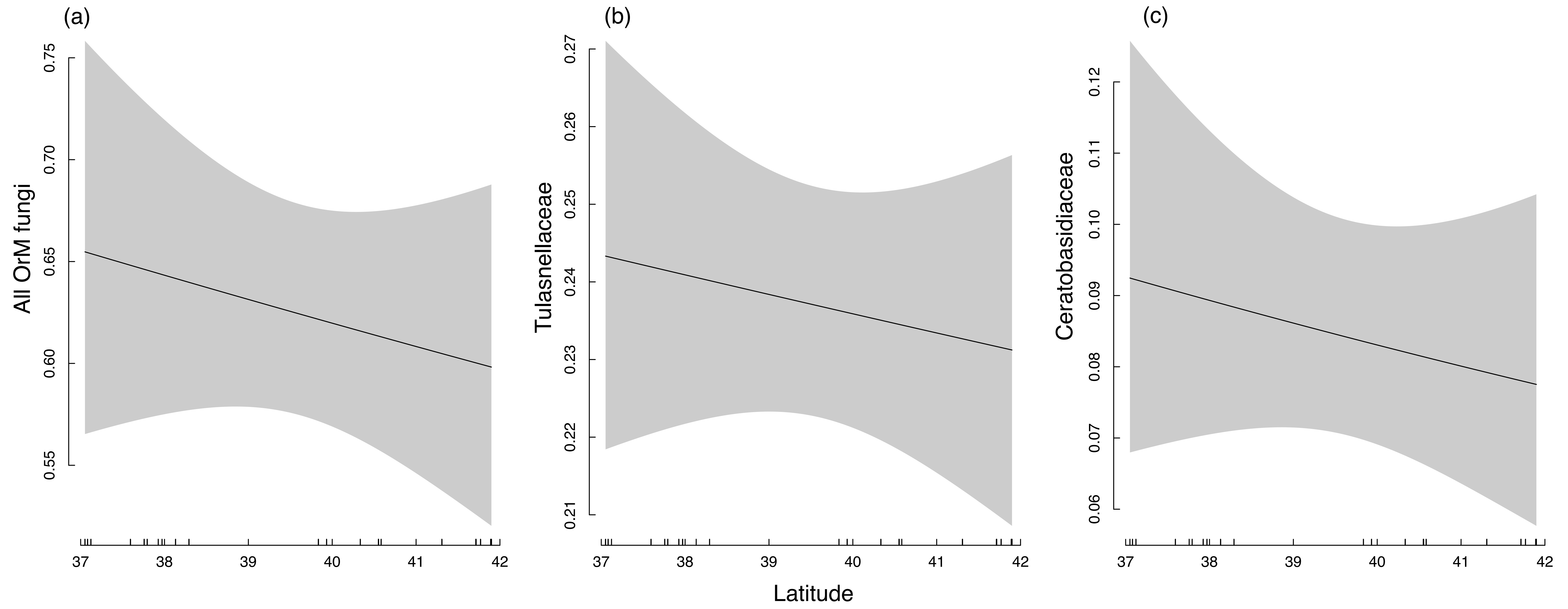

### Fig. S9

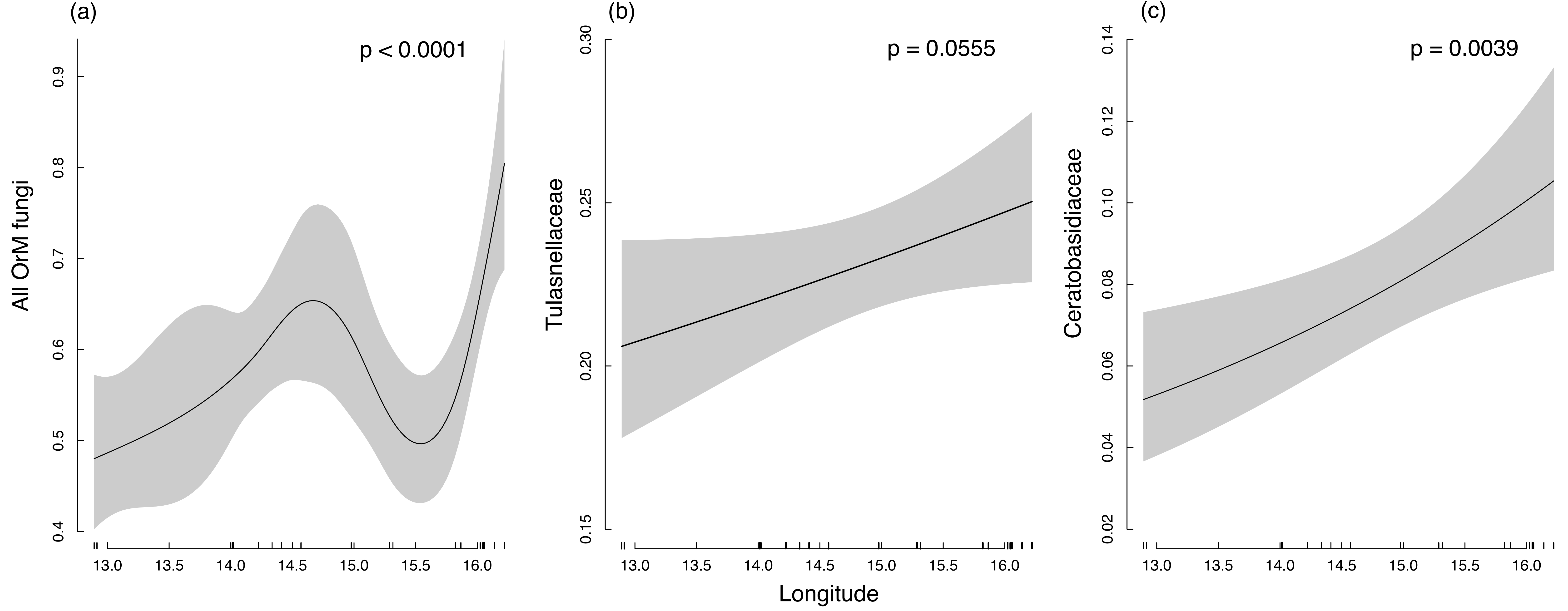

### Fig. S10

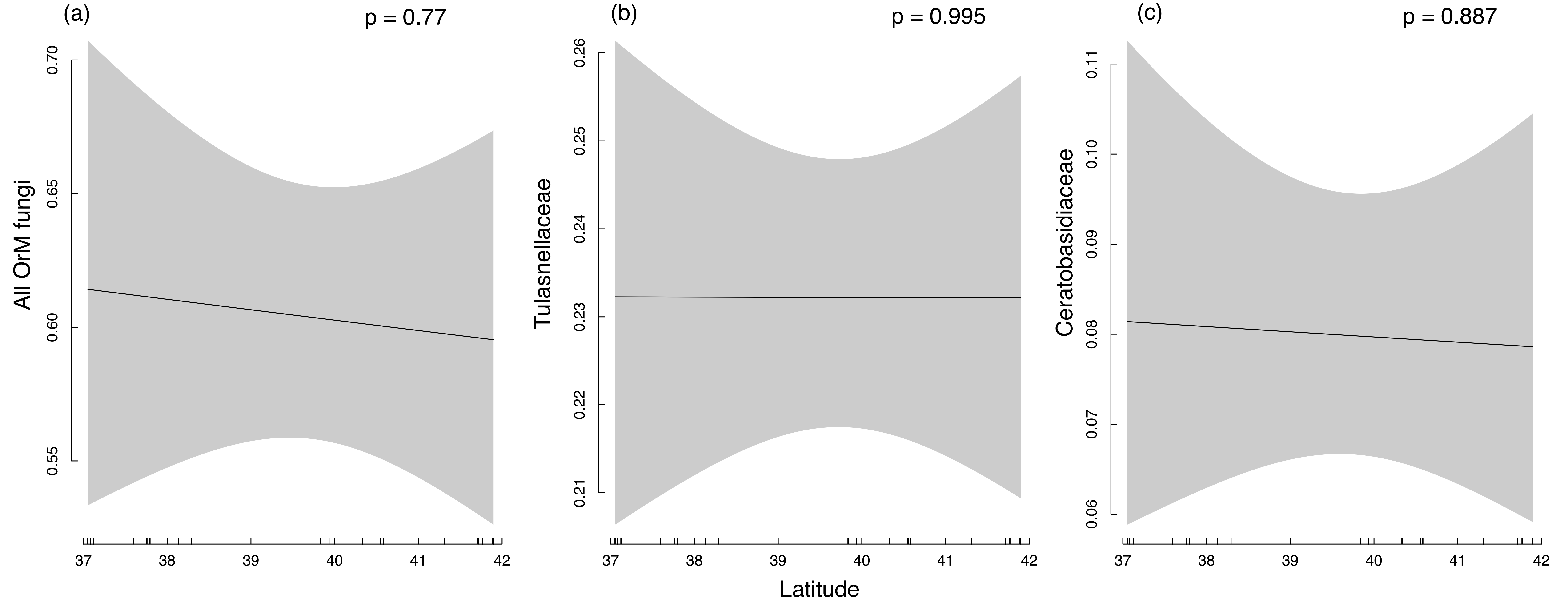

### Fig. S11

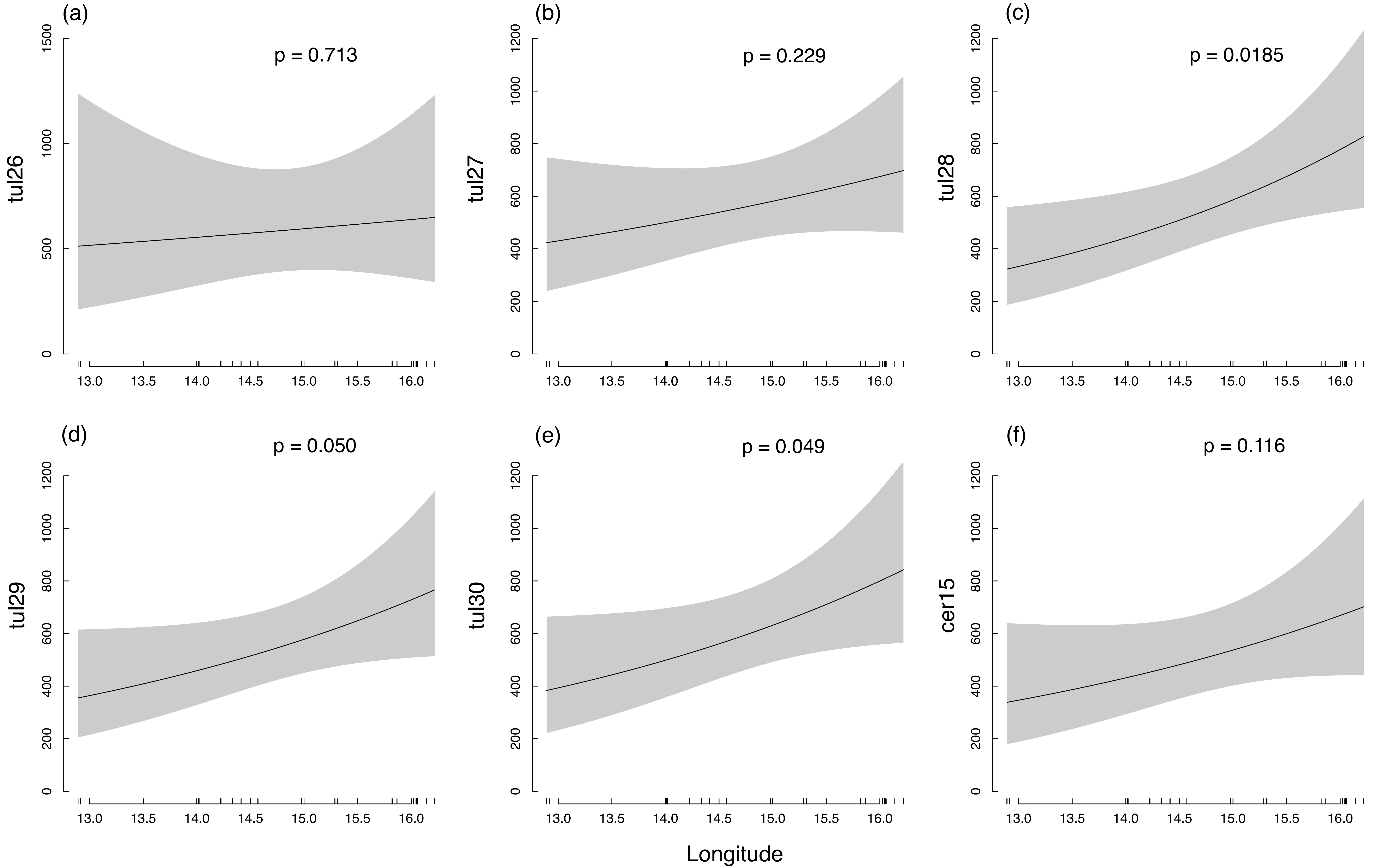

### Fig. S12

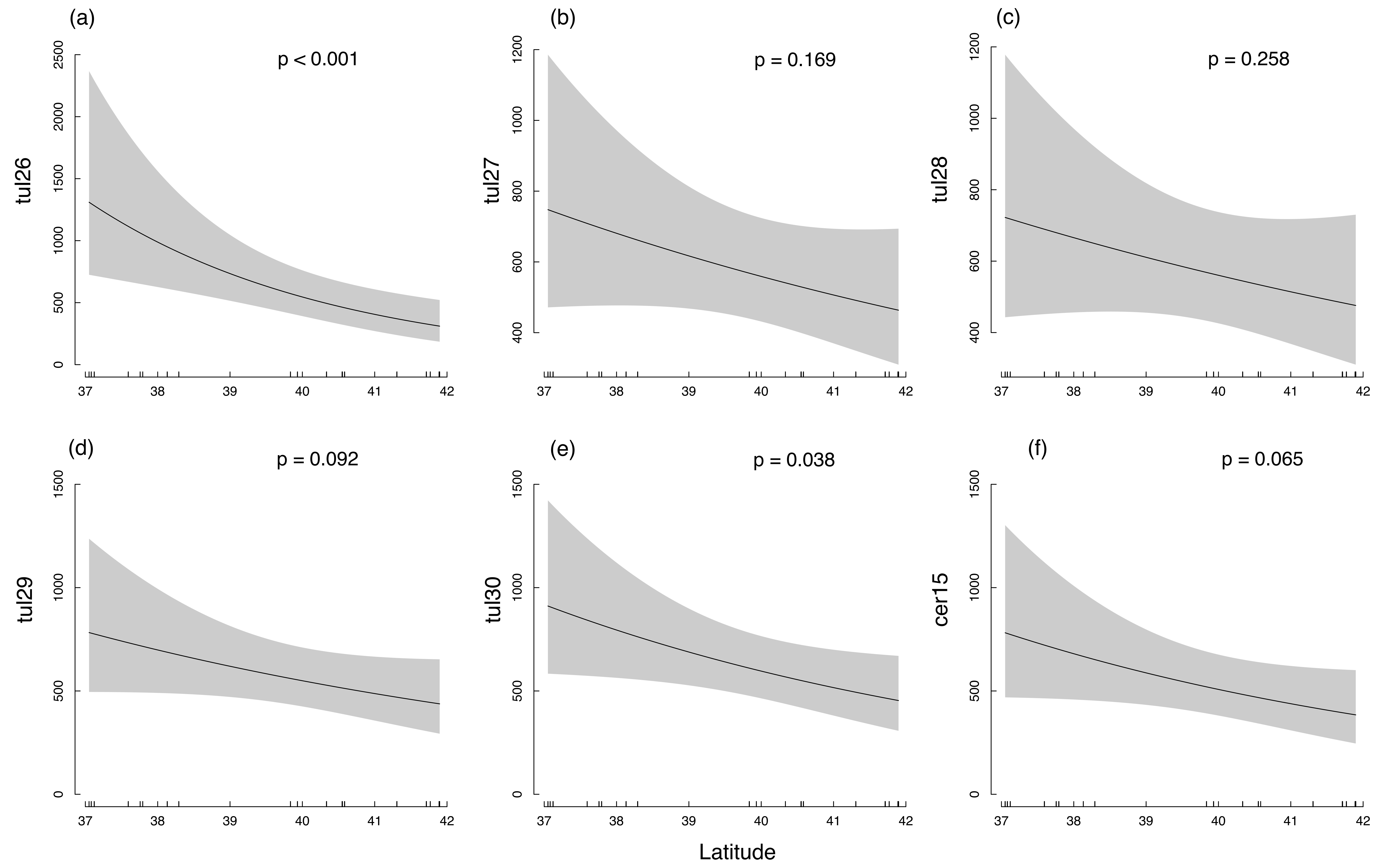
